# Genomic analysis of dingoes identifies genomic regions under reversible selection during domestication and feralization

**DOI:** 10.1101/472084

**Authors:** Shao-jie Zhang, Guo-Dong Wang, Pengcheng Ma, Liang-liang Zhang, Ting-Ting Yin, Yan-hu Liu, Newton O. Otecko, Meng Wang, Ya-ping Ma, Lu Wang, Bingyu Mao, Peter Savolainen, Ya-ping Zhang

**Author notes:** S.J.Z., G.D.W., and P.M. contributed equally to this work.

## Abstract

Dingoes *(Canis dingo)* are wild canids living in Australia. They have lived isolated from both the wild and the domestic ancestor and are a unique model for studying feralization, the process in which a domestic species escapes human control, adapts to the wild, and diverges from the domestic ancestor into a genetically distinct population. Here, we sequenced the genomes of 10 dingoes and 2 New Guinea Singing Dogs, to study the origins and feralization process of the dingo. Phylogenetic and demographic analyses show that dingoes originate from domestic dogs in southern East Asia, which migrated via Island Southeast Asia to reach Australia 4300-5000 years ago, and subsequently diverged into a genetically distinct population. Selection analysis identified 99 positively selected genes enriched in starch and fat metabolism pathways, indicating a diet change during feralization of dingoes. Interestingly, we found that 14 genes have shifted allele frequencies compared to dogs but not compared to wolves. This suggests that the selection affecting these genes during domestication of the wolf was reversed in the feralization process. One of these genes, *ARHGEF7,* may promote the formation of neural spine and synapses in hippocampal neurons. Functional assays showed that an A to G mutation in *ARHGEF7,* located in a transcription factor-binding site, decreases the endogenous expression. This suggests that *ARHGEF7* may have been under selection for behavioral adaptations related to the transitions in environment both from wild to domestic and from domestic back to wild. Our results indicate that adaptation to domestication and feralization primarily affected different genomic regions, but that some genes, related to neurodevelopment, metabolism and reproduction, may have been reversibly affected in the two processes.

## Introduction

Domestication is the process when a wild species is bred in captivity and modified by artificial selection, becoming phenotypically and genetically distinct from the wild ancestor^1, 2, 3, 4, 5, 6^. Feralization is, in a sense, the reverse process, when a domestic species escapes human control, adapts to the wild through natural selection, and diverges from the domestic ancestor into a genetically distinct population^7^. In the shift from artificial to natural selection, feralization is accompanied by phenotypic changes resulting in a phenotype closer to that of the original wild ancestor than to the domestic type. For instance, feralized rodents tend to look more like wild than domestic rodents^8^, feral chicken on the island Kauai has increased brooding like wild Red Junglefowl^9^, and the dingo’s hunting and social behavior is more similar to that of the wolf than of the dog^10, 11, 12^. In plants, weedy rice (a feralized rice population) has a closer semblance to wild than to domestic rice for several growth characters^13, 14, 15^.

Although the feralization process has aroused considerable research interest, only limited research about the genomic mechanisms involved in this phenomenon has so far been presented. A major obstacle for such studies is that, in most cases, the feral populations are not isolated from the wild and/or domestic ancestors, implying a problem to distinguish genetic change caused by feralization from change caused by crossbreeding with the ancestral populations. So far, only two comprehensive studies of genomic changes under feralization have been performed, on feral chicken on Kauai, and on Chinese weedy rice^9, 14^. The research on feral chicken showed adaptation of genes associated with sexual selection and reproduction but suggested that feralization and domestication mostly target different genomic regions ^9^. Similar conclusions were reached concerning the feralization of Chinese weedy rice ^14^, suggesting convergent evolution of different weedy types but little overlap of genes under selection in the domestication and feralization processes. However, both these studies have problems to distinguish genetic change caused by feralization from change caused by crossbreeding with the ancestral populations.

The dingo *(Canis dingo)* is a wild canid native to Australia, and its apex predator ^16^. It originates from domestic dogs but has, since it arrived at least 3,500 years ago, developed into a phenotypically and genetically distinct population of feral dogs. Its appearance is similar to the domesticated dog but there are big differences in its behavior^17, 18^. Like the wolf, the dingo is a predominantly meat-eating omnivorous animal, and lacks the duplications of the alpha-amylase locus giving improved starch digestion in domestic dogs^11, 19, 20^. Dingoes hunt in the wild, can catch and kill large prey such as kangaroos, cattle, water buffalos, and wild horses, and use the same tactics as their wild ancestor, wolves, to hunt the large prey ^10^. While young dingoes are often solitary, adults often form a settled group, and the dingo’s social behavior is as flexible as that of a coyote or gray wolf^12, 21^. The dingo is an ideal and unique model for studying the evolutionary and genomic mechanism of feralization, because of two features. Firstly, the dingo population has a longer history of feralization than any other animal, since their arrival in Australia at least 3500 years ago^22, 23^. Secondly, the dingoes have been isolated from both their domestic and their wild ancestor during this feralization, because of Australia’s position outside the natural range of wolves and its isolation until the arrival of Europeans. Therefore, unlike the feral chickens of Kauai island and weedy rice, the dingoes have not experienced hybridization with ancestral populations which may complicate the deciphering of the genomic mechanisms of feralization.

In the present study, we sequenced the nuclear genomes of 10 dingoes from across Australia and 2 New Guinea Singing Dogs (NGSDs; wild canids living in highland New Guinea), and retrieved the genomes of a worldwide representation of 78 dogs and 21 wolves from literature. Based on this, we analyzed population structure and phylogenetic structure to assess the detailed demographic history and migration route of dingoes, and we performed selection scans to decipher the genomic mechanism of natural selection under feralization and to reveal the correlation between selection in domestication and feralization.

## Results

### Sample collection and whole genome sequencing

10 dingoes and 2 NGSDs were sequenced for the current study. The samples of dingoes have a wide distribution across Australia (Fig. 1A), and the two NGSDs are from the NGSD Conservation Society stud book. After DNA extraction, individual genomes were sequenced to an average of 14.7×. We also retrieved 97 canine whole-genome sequences from published articles^3, 20, 24, 25, 26^, which involved 1 dingo, 1 Taiwan village dog, 43 indigenous dogs from China and Vietnam, 19 individuals from various breeds, 4 village dogs from Africa, 6 Indian village dogs, 3 village dogs from Indonesia, 3 village dogs from Papua New Guinea, and 21 wolves from across Eurasia (Supplementary table 1). Downloaded data have a high quality and an average sequencing depth of 14.6×. Overall, the dataset covers all major dog and wolf groups^27^ that are putative ancestors of dingoes. Raw sequence reads were mapped to the dog reference genome (Canfam3) using the Burrows-Wheeler Aligner (BWA)^28^. DNA sequence analysis was done using the Genome Analysis Toolkit^29^ (see materials and methods). After strict filtering, we identified ~24.7 million autosomal SNPs for further analysis.

**Fig. 1.**
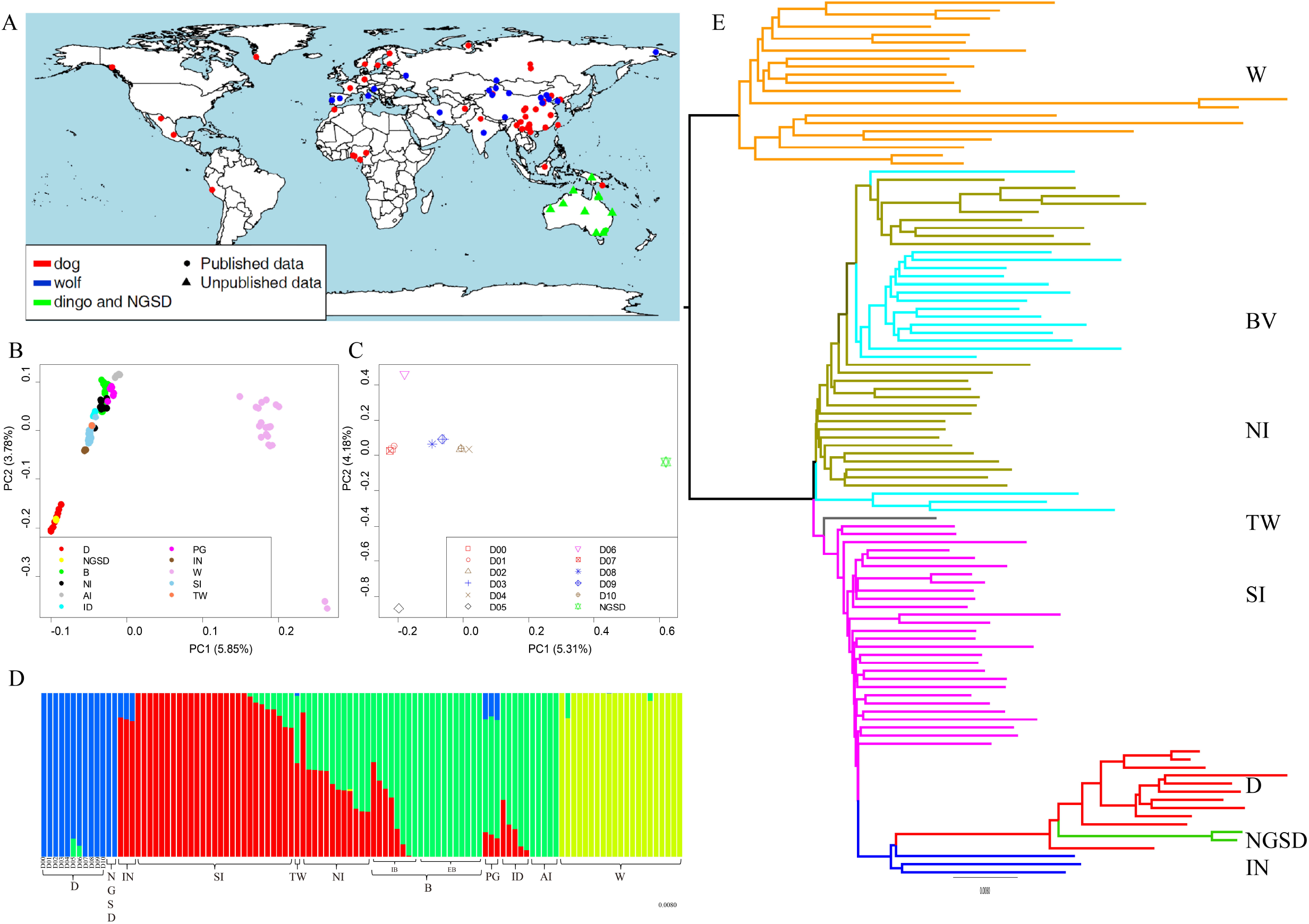
Population structure and genetic diversity of 109 canids. A) Geographic locations of the 109 canids analyzed in this study. B) Principle component analysis of the 109 canids. C) Principle component analysis of only the dingo and NGSD, red = dingo from the Southeast Australia; blue = dingo from West/central Australia; brown = dingo from Northeast Australia; green = NGSD. D) Structure analysis of all the 109 individuals. E) A phylogenetic tree for all the 109 individuals. D, dingoes; NGSD, New Guinea Singing Dogs; IN, Indonesian village dogs; SI, indigenous dogs from southern China; TW, Taiwan indigenous dog; NI, indigenous dogs from north China; B, breeds; IB, intermediate breeds; EB, European breeds; PG, Papua New Guinea village dogs; ID, Indian village dogs; AI, African village dogs; W, wolves; BV, breeds and the other regions village dogs.

### Population structure and phylogenetic analysis

Principal component analysis (PCA) of the 109 individuals was performed to explore the relationships among dingoes, NGSDs, and other canids. In a two-dimensional plot of the genotypes, there is a clear separation in four groups: Tibetan wolves, other wolves, dogs and dingoes/NGSDs. Dogs can be divided into two basic groups: dogs from Europe and indigenous dogs from Asia. All dingoes and NGSDs cluster together tightly, on a relatively large distance from the dogs. Thus, the dingo and NGSD populations are genetically clearly distinct from domestic dogs. Among the dogs, Indonesian village dogs cluster closest to the dingoes/NGSDs, followed by indigenous dogs from southern East Asia (Fig. 1B). Notably, India and Taiwan have been suggested as possible origins for the dingo^30, 31^, but the dogs from these regions cluster relatively far from the dingoes. We then analysed the 10 dingoes and 2 NGSDs separately, to explore their detailed structure (Fig. 1C). The two-dimensional plot separates NGSDs from the dingoes. The dingoes cluster together (except the two individuals D05 and D06, see below; structure analysis indicates hybridization with European dogs), but are distributed in three sub-clusters in accordance with geographical origin: Southeast, West/central and Northeast Australia. This suggests that there are subpopulations within the dingo population.

To explore the genetic relationships among the 109 individuals, we also performed a structure analysis using expectation maximization (EM) algorithm to cluster the individuals into different numbers of groupings. Partitioning the individuals into four groups separated the samples into: i) wolves, ii) dingoes and NGSDs, iii) indigenous dogs from southern East Asia and Indonesia, and iv) breeds and village dogs from other regions (Fig. 1D). We find an admixture of these components with varying proportions among indigenous dogs from northern China, some breeds and the village dogs from India and New Guinea, consistent with the results of the PCA. Notably, two dingoes, D05 and D06, show a mixture indicating hybridization with European breed dogs, and they originate from the two regions in Australia with highest incident of dingo-dog hybridization (the Southeast and the Northwest)^32, 33^. Thus, we performed D-statistics analysis^34^ to test gene flow between the dingo population and European breeds. The results indicate gene flow between the two dingoes (D05 and D06) and European breeds (D=0.1027 and 0.0827, Z=14.551 and 10.221) (Supplementary table 2).

We further performed phylogenic analysis by the Neighbor-Joining (NJ) approach (Fig. 1E and supplementary figure 1). The result matches the observations from the PCA and structure analysis. First, dogs and dingoes/NGSDs separate from the wolves, and then they further split into two clades, one including dingoes and NGSDs together with indigenous dogs from Indonesia and southern East Asia, and the other including village dogs and breeds from all other regions. Indonesian village dogs are closest to the dingoes and NGSDs, and dogs from southern East Asia are basal to the clade. This suggests that indigenous dogs from southern East Asia may be the ancestors of dingoes. Notably, similarly to the PCA, the dogs from India and Taiwan cluster in the second clade, far from the dingoes. We also made a Maximum-Likelihood phylogenetic tree of all the 109 individuals and a NJ phylogenetic tree for 107 individuals (removing D05 and D06, because of the suspected hybridization with European dogs) (Supplementary figures 2-3), obtaining consistent results. Notably, the branch to dingoes and NGSDs is relatively long. We therefore estimated nuclear diversity using the parameter θπ, grouping individuals into four populations (wolves, indigenous dogs from southern East Asia, European breeds, and dingoes (Supplementary figure 4). The result shows that dingoes had the lowest diversity of the four populations (Supplementary figure 5), mean value: 6.94×10^−4^, compared to 1.24×10^−3^ and 1.37×10^−3^ for European breeds and southern East Asian indigenous dogs, respectively. This suggests a severe bottleneck event in the evolutionary history of dingoes, which may explain the long phylogenetic branch.

### Demographic and migration histories

Based on the PCA and phylogeny analysis, the indigenous dogs from southern East Asia are plausible ancestors of dingoes and NGSDs, with the Indonesian village dogs as the most closely related population. To study the migration and demographic history of the dingoes, we performed a demographic analysis using G-PhoCS^35^. Based on a mutation rate of 2.2 × 10^−9^ per site per year^36^, our analysis indicates that the split between dingoes and Indonesian village dogs occurred around 4,900 years ago and that, previous to that, Indonesian village dogs and the indigenous dogs from southern East Asia diverged around 5,800 years ago (Fig.2A; Supplementary table 4). It also shows that the dingo population has a very small effective population size, and it indicates gene flow from southern East Asia to Indonesian village dogs. We then used smc++, a software employing unphased whole genomes to infer population history^37^. This analysis approximated the split between Indonesian village dogs and dingoes at around 5,400 years ago, consistent with the result of G-PhoCS. We also used smc++ to estimate dates for the population history of dingoes/NGSDs and dogs. The result shows that the dog population experienced a slight growth after the population split, while the dingo and NGSD populations suffered a decrease (Fig.2B), followed by an increase possibly reflecting the expansion into the new ecological niches in Australia and New Guinea. Notably, the NGSDs show a severe decrease in more recent times followed by a sharp increase. This is consistent with the history of the western population of NGSDs (bred outside New Guinea the last 60 years), which originates from very few individuals.

### Mitochondrial genome analysis

We also performed phylogenetic analysis based on mitochondrial genomes, analyzing totally 35 dingoes and 3 NGSDs, the 10 dingoes and 2 NGSDs sequenced in this study and 25 dingoes and 1 NGSD from Cairns et al.^38^, in the context of 169 dogs and 8 wolves from across the Old World from Pang et al.^39^. We constructed a phylogenetic tree showing all dingoes and NGSDs to group into a single branch, separated from all domestic dogs except one, a dog originating from Hunan in South China (Fig. 3A). The dingo/NGSD branch is part of the major domestic dog haplogroup A, to which approximately 75% of domestic dogs worldwide belong. Haplogroup A has six sub-haplogroups, and the dingo/NGSD branch is part of sub-haplogroup A2, which is frequent in dogs originating from across East Asia but absent in western Eurasia^39^. Notably, of the eight dogs clustering closest to the dingo/NGSD branch, seven were from Mainland or Island Southeast Asia and one from East Siberia. These results indicate that dingoes and NGSDs originate from domestic dogs in Southeast Asia, via Island Southeast Asia, and that dingoes and NGSDs are closely related, as earlier suggested^22, 38, 40, 41^.

**Fig. 2.**
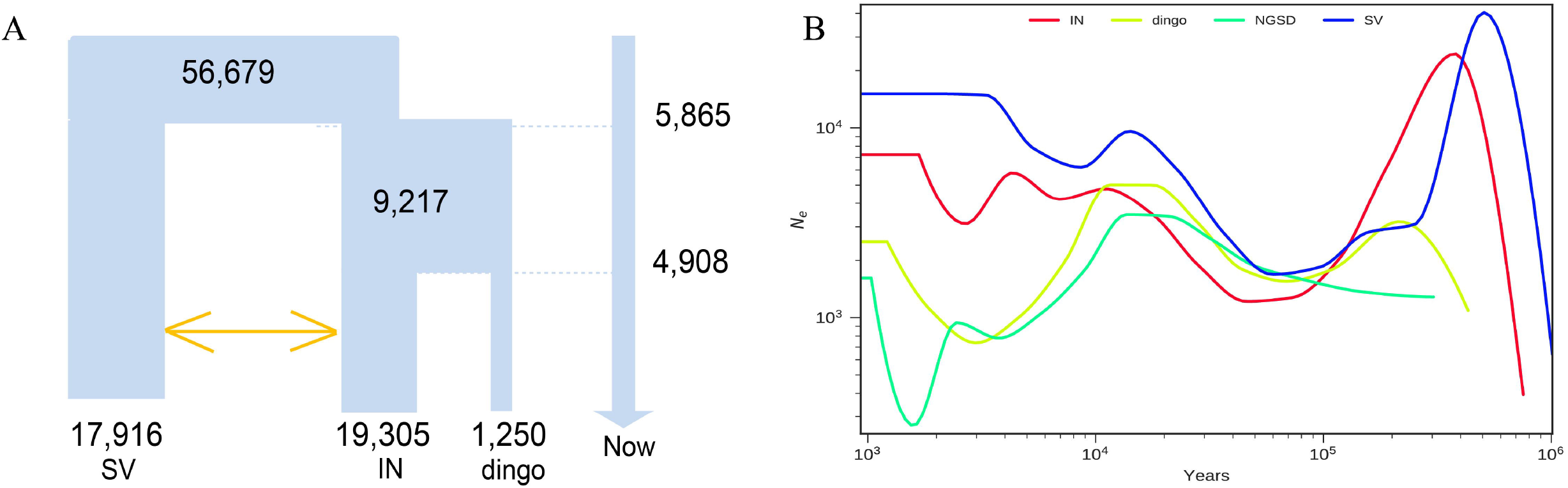
Demographic history of dingo and other dogs. A) Demographic history inferred for indigenous dogs from southern China (SV), Indonesian village dog (IN) and dingo using G-phocs. B) Inferred effective population sizes with the time for indigenous dogs from southern China (SV), Indonesian village dog (IN), dingo and NGSD using SMC++.

**Fig. 3.**
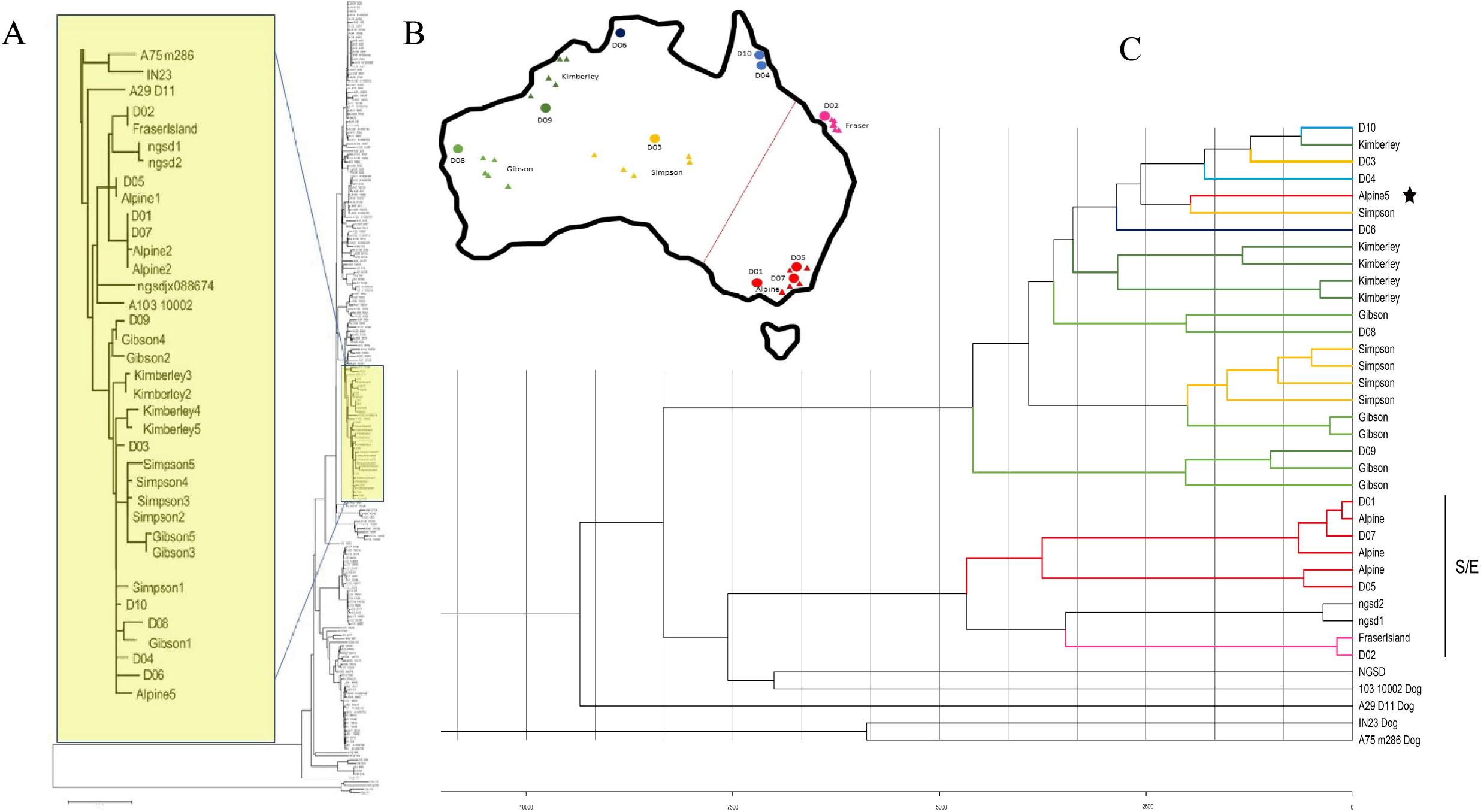
Phylogenetic and demographic history analysis of mtDNA. A) Neighbor joining tree based on mitochondrial genomes from 35 dingoes and 3 NGSDs, and from 169 domestic dogs and 8 wolves from across the Old World, with 4 coyotes as outgroup. The yellow box and inset figure indicates the branch in which all dingoes and NGSDs cluster together with a single domestic dog from South China, and the 3 most closely related dogs outside this branch. B) Map depicting geographic sampling of dingoes across Australia. Circles represent the 10 individuals sequenced in this study and triangles 25 additional samples from Cairns et al. The red line indicates the genetic subdivision between the southeastern/eastern part (S/E) and the rest of the continent. C) Bayesian analysis of mitochondrial genomes for the sub-dataset identified in Fig. 3. The dingo/NGSD branch including all dingoes and NGSDs and a single South Chinese domestic dog and, as outgroup, the three most closely related dogs outside that branch. The scale axis indicates time estimates using the mutation rate of 7.7× 10-8 per site per year with SD 5.48×10-9 from Thalmann et al. The colored branches indicates geographical origin of dingo samples, see Fig. 2. The star highlights the single dingo sample from southeast Australia that does not cluster in the S/E-related branch.

To study the detailed phylogeny among dingoes and NGSDs we created a sub-dataset including all individuals in the dingo/NGSD branch and the three most closely related dogs (yellow box in Fig.3A), and constructed new phylogenetic trees (Fig. 3C; supplementary figures 6-7). These trees show a division of dingoes into two main branches, following a geographical distribution earlier reported by Cairns et al.^38^; all dingoes from Southeast and East Australia (we denote this region S/E), except one, group in one branch, and all dingoes from all other parts of Australia group in the other (Fig. 3B and 3C). The S/E-related branch also includes two of three NGSD samples, while the third NGSD and the domestic dog from South China have an intermediate position. Notably, outside the S/E region there is only limited geographical structure among the dingoes. Thus, there is a genetic subdivision of dingoes between the southeastern/eastern part of Australia and the rest of the continent, possibly reflecting two (parallel or consecutive) introductions into Australia from an ancestral population in Island Southeast Asia.

Molecular clock analysis (based on a mutation rate of 7.7×10^−8^ per site per year)^42^ suggests a most recent common ancestor (MRCA) for all dingoes and NGSDs (the division into the two main branches) ~8,000 years ago, and MRCAs for each of the two branches ~4,500 years ago. This suggests that both dingo branches were introduced to Australia ~4,500 years ago, in agreement with the nuclear genome estimate for the split between dingoes and Indonesian dogs.

### Natural selection in feralization

Our analyses of population structure and demography confirms that the dingoes originated by feralization of domestic dogs around 5,000 years ago and have since then remained virtually isolated from both the wild and the domestic ancestor until recent time. This affirms that the dingo is an excellent model for studies of the genomic effects of feralization. We used analysis of Fst and iHS ^43^ to identify positive selection in the dingoes. Firstly, pairwise Fst statistics were calculated between dingoes and dogs from southern East Asia (Fst (ds)), using Weir and Cockerham’s method ^44^ implemented in VCFtools v0.1.11 ^45^ with non-overlapping 20 kb genomic windows. With a threshold of Z(Fst)> 2.0, we identified divergent regions spanning totally 4.15% of the autosomal genome. Furthermore, we performed a windowed iHS test ^46^ with the same non-overlapping 20 kb windows. A window was identified if a 0.3 fraction of iHSs had an absolute value above 2, indicating 3.02% of the autosomal genome to be under selection. The overlap of the two approaches indicated 174 candidate windows under positive selection, containing 99 genes (Supplementary table 3) considered as candidates associated with feralization of dingoes. GO enrichment evaluations identified 16 functional classes that were significantly overrepresented (p < 0.05) (Table 1). Groups of genes showing the strongest evidence of positive selection are those related to cellular component assembly, cellular component biogenesis, organelle organization, cytoplasmic part and metabolism.

**Table 1.** Gene ontology analysis of the the 99 feralization gene candidates

| Category | GO Term | Num. of Genes | P-Value |
| --- | --- | --- | --- |
| BP | cellular component assembly | 17 | 0.0081 |
| BP | mitotic sister chromatid segregation | 4 | 0.015 |
| BP | mitotic nuclear division | 5 | 0.017 |
| BP | cellular component biogenesis | 17 | 0.02 |
| BP | sister chromatid segregation | 4 | 0.023 |
| BP | cell-substrate adhesion | 5 | 0.025 |
| BP | cell-matrix adhesion | 4 | 0.029 |
| BP | organelle organization | 20 | 0.044 |
| BP | spindle assembly | 3 | 0.047 |
| CC | cytoplasmic part | 34 | 0.043 |
| MF | alpha-1,4-glucosidase activity | 2 | 0.014 |
| MF | alpha-glucosidase activity | 2 | 0.019 |
| MF | 3-hydroxyacyl-CoA dehydrogenase activity | 2 | 0.037 |
| MF | glucosidase activity | 2 | 0.042 |
| KEGG PATHWAY | Fatty acid degradation | 3 | 0.016 |
| KEGG PATHWAY | Fatty acid metabolism | 3 | 0.021 |

Most interestingly, five genes, distributed among six GO terms, have functions related to digestion and metabolism, they may play roles in diet change of dingoes: alpha-1,4-glucosidase activity (GO: 0004558), alpha-glucosidase activity (GO: 0090599), 3-hydroxyacyl-CoA dehydrogenase activity (GO:0003857), glucosidase activity (GO:0015926), and two KEGG pathway (Table 1): Fatty acid degradation (map00071) and Fatty acid metabolism (map01212). And other interesting genes may play roles in feralization adaptation of dingoes: *ACSL6* (Acyl-CoA Synthetase Long Chain Family Member 6) regulates lipid synthesis in human and rat skeletal muscle ^47^, *EHHADH* (Enoyl-CoA Hydratase And 3-Hydroxyacyl CoA Dehydrogenase) is part of the classical peroxisomal fatty acid β-oxidation pathway and essential for the production of medium-chain dicarboxylic acids^48^, and *HADHB* (Hydroxyacyl-CoA Dehydrogenase Trifunctional Multienzyme Complex Subunit Beta) encoded proteins catalyze the last three steps of mitochondrial beta-oxidation of long chain fatty acids. The candidate genes include also an important gene associated with reproduction, *Prss37* (Protease, Serine 37), shown to be required for male fertility in mice ^49^.

### Genes with reversible selection in domestication and feralization

To compare the positive selection in the domestication and feralization steps, we also calculated Fst between wolves and dogs from southern East Asia (Fst (ws)), and Fst between wolves and dingoes (Fst(dw)), in non-overlapping 20 kb windows. The mean values of Fst (dw), Fst (ds), and Fst (ws) were 0.1226, 0.0966, and 0.0766, respectively (Supplementary figure 8) and 67.7% of the 20 kb windows had Fst (dw)> Fst (ds), in agreement with the closer relationship of dingoes to domestic dogs than to wolves.

Importantly, for 14 of the candidate genes, Fst(ws) was significantly high (Z(Fst(ws))>2) while Fst(dw) was relatively low (Z(Fst(dw))<0) (Fig.4), signifying a small difference between wolf and dingo, but a large difference between dog and dingo and between dog and wolf. This suggests that these 14 genes evolved during both the feralization and the domestication steps, such that feralization imposed reversed selection compared to the domestication. Functional annotation showed that four of these 14 genes are associated with neurodevelopment, metabolism and reproduction (Table 2). *ARHGEF7* (Rho Guanine Nucleotide Exchange Factor 7) may promote the formation of neural spine and synapses in hippocampal neurons^50^ and *BPTF* (Bromodomain PHD Finger Transcription Factor) promotes posterior neuroectodermal fate^51^, while *SI* (Sucrase-Isomaltase) has been shown to play an important role in the digestion of malted dietary oligosaccharides and to play important functions in the final stage of carbohydrate digestion^52^, and finally, *Prss37* (Protease, Serine 37) is related to reproduction.

**Table 2.** The 14 genes under selection in both domestication and feralization

| Chr | Start | End | Z(Fst)-ds | Z(Fst)-ws | Z(Fst)-dw | Gene |
| --- | --- | --- | --- | --- | --- | --- |
| 1 | 99580001 | 99600000 | 3.451028 | 2.394955 | -1.79066 | ENSCAFG000000023509 |
| 5 | 77260001 | 77280000 | 2.902392 | 3.811514 | -0.10461 | ENSCAFG000000029432 |
| 6 | 52100001 | 52120000 | 5.065877 | 2.224988 | -0.57871 | DPYD |
| 9 | 12620001 | 12700000 | 3.386893 | 4.109578 | -2.26201 | BPTF |
| 16 | 7340001 | 7360000 | 3.363463 | 2.644232 | -1.8555 | PRSS37/<br>ENSCAFG00000003879 |
| 18 | 42840001 | 42860000 | 2.845394 | 2.285565 | -1.21919 | ATG13 |
| 19 | 33820001 | 33840000 | 4.113953 | 3.324179 | -0.56627 | DPP10 |
| 22 | 59220001 | 59260000 | 2.367673 | 3.076211 | -0.08149 | ARHGEF7 |
| 30 | 16480001 | 16500000 | 2.53002 | 2.537018 | -0.85537 | TRPM7 |
| 30 | 16540001 | 16580000 | 4.021162 | 6.10273 | -0.83127 | TRPM7 |
| 31 | 39500001 | 39520000 | 2.512109 | 3.886574 | -1.25248 | ENSCAFG000000012030/<br>LSS/MCM3AP |
| 34 | 30360001 | 30380000 | 2.644394 | 2.896281 | -0.037351 | SI |

### Functional assay revealed that a mutation in dingo gives decreased enhancer activity for *ARHGEF7*

*ARHGEF7* is related to neural function ^50^ and may therefore be involved in behavior changes in the development from dog to dingo. We found an A-to-G mutation (chr 22:59233505) within the *ARHGEF7* gene, which had a very high allele frequency in dingoes (100%) and wolves (93.3%) compared to indigenous dogs from southern East Asia (32.5%). Furthermore, detailed bioinformatics analysis showed that the A-to-G mutation may influence the expression of *ARHGEF7* since it is located in a transcription factor-binding site ^53^. To test whether the SNP variants can actually affect expression of *ARHGEF7,* we performed dual-luciferase reporter experiments (enhancer assay using pGL3-promoter vectors) on two human cell lines (Daoy, human medullablastoma and HEK293, human embryonic kidney) and one canine cell line (MDCK, Madin-Darby Canine kidney). The analyses showed that all three cell lines displayed significantly lower enhancer activities for SNP-G than for SNP-A (Fig. 5), suggesting that SNP-G may confer decreased endogenous *ARHGEF7* production.

**Fig. 4.**
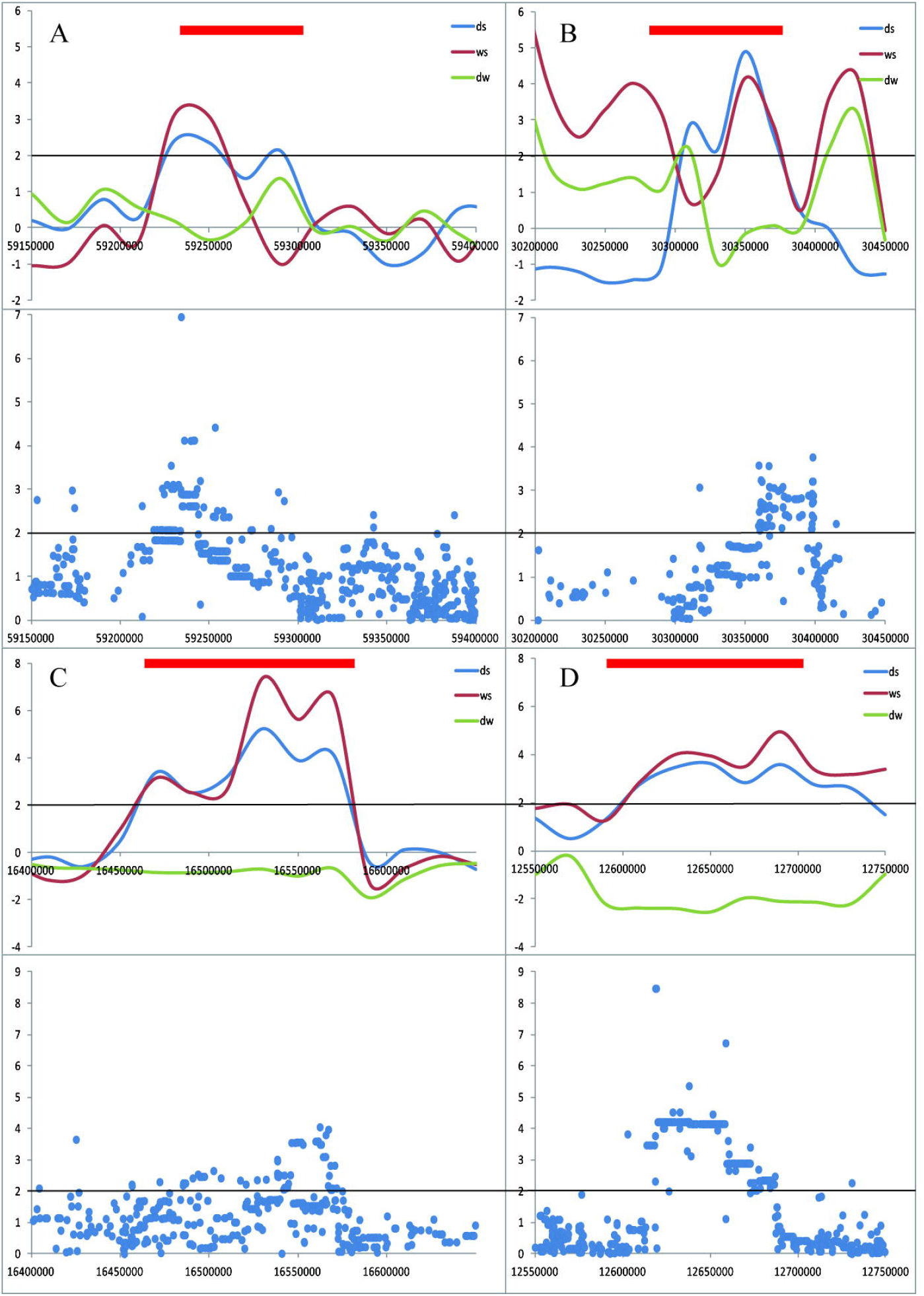
Selection and allele-frequency differentiation between feralization and domestication. Fst between dogs from southern East Asia and dingo, denoted “ds”, blue; Fst between wolf and dogs from southern East Asia, denoted “ws”, red; Fst between wolf and dingo, denoted “dw”, green. A) ARHGEF7, B) SI, C) TRPM7, D) BPTF. Thick red horizontal line indicates range of gene.

**Fig. 5.**
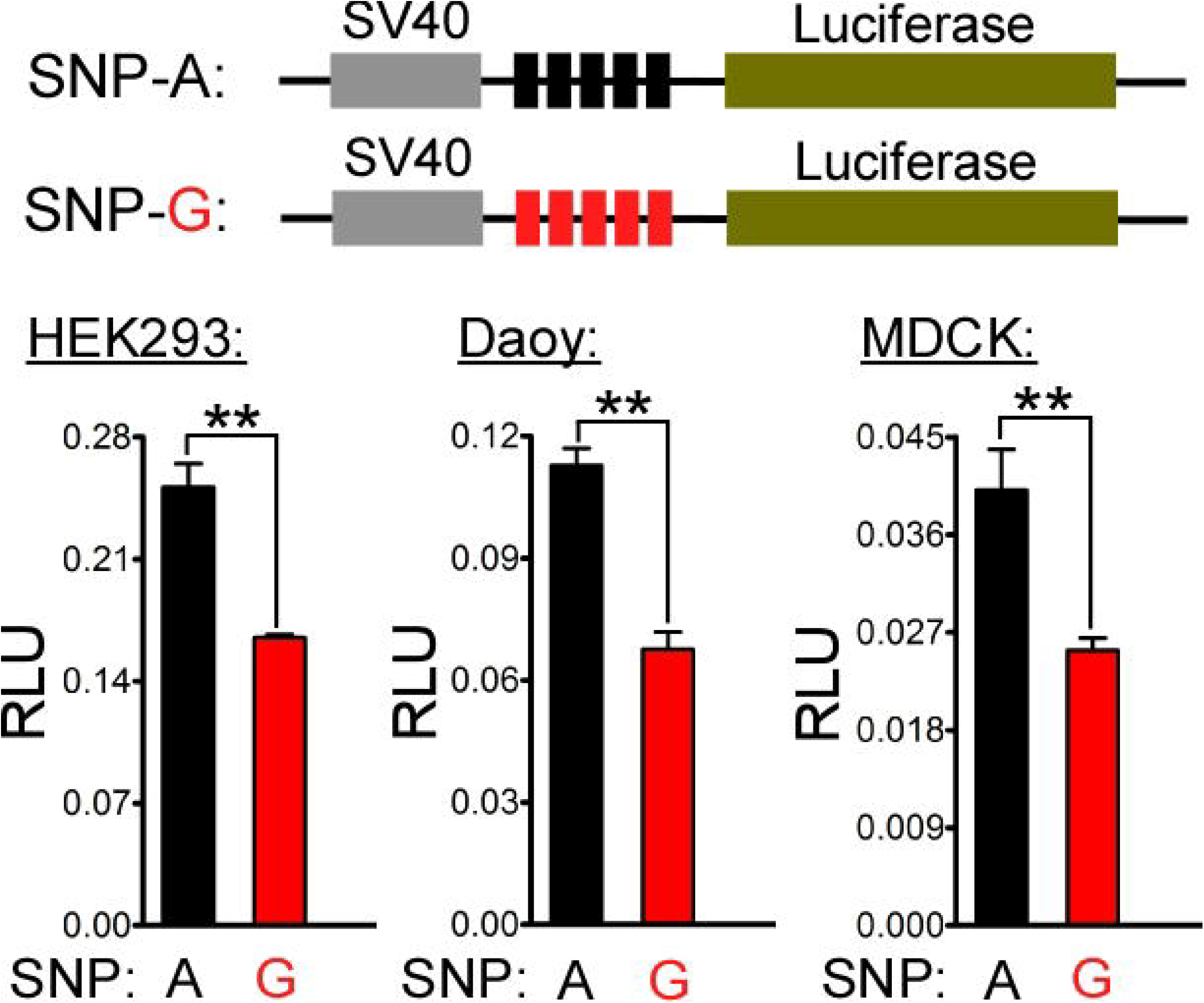
Functional assay of ARHGEF7. Dual-luciferase reporter experiments (the enhancer assay using pGL3-promoter vectors) using two different human cell lines (HEK293, human embryonic kidney and Daoy, human medullablastoma) and one canine cell line (MDCK, Madin-Darby Canine kidney).

## Discussion

In this work, we have investigated the process of feralization on the genomic level, using the dingo as a model. The analyses of population structure and demography reinforces that the dingo is an excellent model for this, because its feralization started 4,000-5,000 years ago and because it has since then remained isolated from its domestic and wild ancestors. This makes the dingo a unique tool for identifying genomic regions under positive selection in the feralization process without confusing the impact of feralization with hybridization to ancestral populations.

Our analyses identify 99 candidate genes in genomic regions under selection, and find an overrepresentation of genes correlated in particular to digestion, metabolism and reproduction. This indicates an adaptation to a new environment, in the form of a change of diet and changed sexual and reproductive mechanisms. This agrees with the two previous studies of genomic change under feralization, on feral chicken and rice. In the feral rice, genomic regions containing numerous genes correlated with adaptation to the new environment were identified, linked to, e.g., flowering time, reproduction and stress response^14^. In the feral chickens, especially genes correlated with sexual selection and reproduction were identified, e.g., genes correlated with fecundity traits, which may be targets of selection that facilitated the feralization^9^. It is notable that genes correlated with sexual selection and reproduction were identified in feral chicken and rice as well as in the dingo, indicating that change in reproduction mechanisms is an important effect of feralization in both animals and plants.

There is considerable difference in diet between domestic dogs and the two related wild canids, wolves and dingoes. The wild canids have a diet consisting predominantly of meat, while domestic dogs normally eat considerable amounts of vegetable food, provided by humans^20, 25, 36, 54, 55^. This diet change has been shown to be reflected by strong selection for improved digestion of starch in domestic dogs36. This is manifested most prominently by expansion of copy numbers of the gene for pancreatic amylase *(AMY2B)* in most dogs, but dingoes have the non-expanded wild type found in wolve^56, 57^. In our selection analyses we now also identified three genes coding for proteins involved in digestion and metabolism of fatty acids. This indicates that diet change has implied a major environmental influence on the dingo, resulting in genomic change. Interestingly, the demographic analysis indicates that the dingo ancestors originated from southern China 5,300-6,600 years ago, at which time large-scale rice farming was established in this region^57, 58, 59^.

We also compared the genomic regions under selection in the feralization steps to those selected in the domestication step, by comparing Fst among wolf, dog and dingo. This analysis identified 14 genes in regions under selection in both the domestication step and the feralization step. This indicates that while adaptation to domestication and feralization has primarily affected different genomic regions, there are some regions for which feralization and domestication overlap and feralization is in fact a reverse process of domestication, we call this process as reversible selection. Interestingly, four of the genes are associated with neurodevelopment, metabolism and reproduction. Notably, Dingoes have several traits that are closer to those of wolves than of dogs, e.g., hunting and social behavior, and diet^10, 11, 12^. Our results suggest that some of these traits may be the result of reversed selection during feralization and domestication.

Two of the 14 genes are related to neurodevelopment, and therefore possibly involved in behavior change necessary for feralization. We performed a functional analysis on one of these genes, *ARHGEF7* which promotes the formation of spines and synapses in hippocampal neurons^50^. This test showed that a SNP found in dingo, located in a transcription factor-binding site, gives significantly lower enhancer activities. Hippocampus plays important roles in response inhibition, memory, and spatial cognition^60, 61^, and some studies suggest that hippocampus relates to purposive behaviorism ^62^. Therefore, changes in expression of this gene may be related to behavior changes in the dingo, linked to the adaptations to a wild environment.

Our study has also presented important new findings about the origins and history of the dingo. In the past decades, numerous population genetic studies of the dingo have been performed based on mitochondrial and Y-chromosomal DNA^22, 30, 40, 41^, indicating an origin from East Asian domestic dogs but lacking in precision about timing, routes of arrival to Australia and demographics. Our studies of whole genomes in dingoes and related canids clarifies several of these details. Our analyses of phylogeny, population structure, demography as well as selection analysis show that the dingo is a genetically distinct population clearly differentiated from the domestic dog. The selection analyses indicate that 4,000-5,000 years of feralization has affected numerous genes linked to, e.g., neurodevelopment, metabolism and reproduction.

The genomic data provides strong evidence that the dingo originates from domesticated dogs in southern East Asia, which migrated via Island Southeast Asia 5300-6600 years ago, to eventually reach Australia 4300-5000 years ago. It is notable that the mitochondrial genome data is in good agreement with these findings. With this data, we can reject two previous hypotheses about the origin and migration routes of dingoes. Based on similarity in skeletal anatomy to Indian pariah dogs and wolves, and gene flow from ancient Indians to indigenous Australians dated at BP4200^63^, it has been suggested that the dingo ancestors came from India^23, 31^. An alternative theory has been that dingoes originate from dogs introduced with the Austronesian expansion into Island Southeast Asia, which arrived in New Guinea about 3,600 years ago^22^. However, the genomic analyses, as well as previous mtDNA data, clearly indicate an origin from dogs in southern East Asia, which arrived to Australia via mainland Southeast Asia, and our demographic analysis indicates an arrival in Australia 4300-5000 years ago, well before the Austronesian expansion^64, 65, 66^. Thus, the genetic data clearly suggest that the dingoes originate from domestic dogs in southern East Asia that migrated via mainland and Island Southeast Asia to reach Australia 4300-5000 years ago, but the human population that was involved in this migration remains unknown.

The results showed that dingoes and NGSDs are genetically very closely related, indicating a common origin from dogs in Island Southeast Asia around 5,000 years ago. We also note that there is a phylogeographic structure in the dingo population recorded by nuclear as well as mitochondrial data. This possibly relates to an origin from more than one introduction to Australia, but if so from a very homogenous source population and at similar points of time.

In this study, we demonstrate that the feralization of the dingo induced positive selection on genomic regions correlated to neurodevelopment, metabolism and reproduction. While adaptation to domestication and feralization has primarily affected different genomic regions, there are some regions for which feralization and domestication overlap and feralization acts as a reverse process of domestication. We also establish that the dingo originated 4,300-5,000 years ago from domestic dogs in southern East Asia. The dingo has thereafter remained isolated, and under 4,000-5,000 years of adaptation to a life in the wild it has developed into a genetically distinct population clearly differentiated from its domestic ancestors.

## Materials and Methods

### Samples and sequences

We examined whole-genome sequences from the largest and most diverse group of dingo studied to date, amassing a dataset of 109 canines around the world. We sequenced genomes of 10 dingoes and 2 New Guinea Singing Dogs in the study. Total genomic DNA was extracted from the blood or tissue samples of the animals using the phenol/chloroform method. For each individual, 1-3 μg of DNA was sheared into fragments of 200-800 bp with the Covaris system. DNA fragments were then processed and sequenced using the Illumina HiSeq 2000 platform. Sequence data pre-processing and variant calling. Raw sequence reads were mapped to the dog reference genome (Canfam3)^67^ using the Burrows-Wheeler Aligner (BWA)^28^. Sequence data were next subjected to a strategic procedure for variant calling using the Genome Analysis Tool Kit (GATK)^29^. During base and variant recalibration, a list of known SNPs downloaded from the Ensembl database was used as the training set.

### Genetic diversity and population structure

Genetic diversity was calculated using VCFtools ^45^. Principal component analysis was made using the smartPCA ^68^. The NJ phylogenetic tree was built by MEGA ^69^.

### Population history

We inferred a complete demographic model for dingo and other dogs, including population divergence times, population size using the G-PhoCS^35^. The Generalized Phylogenetic Coalescent Sampler (G-PhoCS) was employed to infer the complete demographic history for dingo, including population divergence times, ancestral population size, and migration rates based on 1000 neutral loci. The parameters were inferred in a Bayesian manner using a Markov Chain Monte Carlo (MCMC) to jointly sample model parameters and genealogies of the input loci. Burn-in and convergence of each run were determined with TRACER 1.5 ^70^. For the control file of G-PhoCS, Divergence times in units of years, effective population sizes, and migration rates were calibrated by the estimates of generation time and neutral mutation rate from previous studies. A generation time of 3 years and a neutral mutation rate of 2.2e-09 per site per year were used to convert the population sizes and scaled time into real sizes and time.

### Mitochondrial genome analysis

The NJ phylogenetic tree was built using MEGA ^69^ Bayesian analysis was made using Beast^71^, assuming a mutation rate of 7.7×10^−8^ per site per year with SD 5.48×10 according to Thalmann et al.^42^ Burn-in and convergence of each run were determined with TRACER 1.5 ^70^.

### Selection analysis

We make Fst by Vcftools between dingoes, SE Asia/South China dogs and gray wolves in 20K window, then we normalized the fst by the method of zero-mean normalization with a threshold of Z(Fst)> 2.0. We make iHS in dingo by the software of selscan, and also normalized the scores by zero-mean normalization. Then, we performed a windowed iHS test, where a 20kb window of the genome and identify candidate regions for selection if 0.3 fraction of iHS had absolute value above 2. At last, we use a method based on fst to search the population genetic differentiation between wild ancestor population, domestication population and feralization population, to find genetic difference impact by the different process of evolution. We use Fst to stand for the changes of gene frequency.

### Functional test using dual-luciferase reporter assay

To construct *ARHGEF7* enhancer SNP reporters, we insert five repeats of the 50bp fragments arounding the indicated SNP site into the pGL3-Promoter vector (Promega) within MluI and XhoI sites. We verified all recombinant clones by sequencing. Daoy (human medullablastoma), HEK293 (human embryonic kidney) and MDCK (Madin-Darby Canine Kidney) cells were cultured in high-glucose Dulbecco’s modified Eagle’s mediun (DMEM) (Corning) with 10% fetal bovine serum (FBS) (Gibco) at 37°C in 5% CO2 condition. For luciferase reporter assays, Daoy, HEK293 and MDCK cells were transfected with the indicated reporter plasmids together with the same TK-Renilla internal control reporter vectors by using the lipofectamine 2000 transfection reagent (Invitrogen) and changed with the fresh medium at 6 hours after transfection. According to the manufacture’s instruction, luciferase activity was measured at 36 hours after transfection by using the Dual-Luciferase Reporter Assay System (Promega). All assays were performed in at least three independent experiments with a minimum of three replicates.

## Acknowledgements

We thank J. William O. Ballard of the University of New South Wales for providing dingo samples and Janice Koler-Matznick for providing NGSD samples. Australian Government Export permit number N39585 and University of New South Wales Ethic’s Approval 16/77B to Professor Bill Ballard. This work was supported by grants from the NSFC (91531303 and 31571353), and the Breakthrough Project of Strategic Priority Program of the Chinese Academy of Sciences (CAS) (XDB13000000). G-D. W. is supported by the Youth Innovation Promotion Association of CAS and the 13th Five-year Informatization Plan of CAS (Grant No. XXH13503-05).

## Accession codes

The raw sequence data from this study have been submitted to the GSA (http://gsa.big.ac.cn/) under accession CRA000200 for raw data of genomes.

